# GARSA: An integrative pipeline for genome wide association studies and polygenic risk score inference in admixed human populations

**DOI:** 10.1101/2023.05.03.539305

**Authors:** Fernando P. N. Rossi, José S. L. Patané, Vinicius de Souza, Jennifer M. Neyra, Rogério S. Rosa, Jose E. Krieger, Samantha K. Teixeira

## Abstract

Genome-wide association studies (GWAS) and polygenic risk scores (PRS) are multistep analytical tools to identify genetic variants and to assess their contribution to phenotypes/diseases. These analyses are evolving and becoming instrumental to understand the genetic architecture of complex phenotypes/diseases. Nevertheless, to date, there is no single solution incorporating all major steps related to those analyses combined with robust populational bias correction. Here, we describe a semi-automated pipeline unifying steps involved in GWAS and PRS including widely used software. Our pipeline handles quality control (QC), GWAS, and PRS steps, managing different types of input/output files. Furthermore, it includes robust bias correction steps, such as inference of kinship matrix with correction for population structure, use of principal component analysis (PCA) with detection and removal of outlier variant followed by re-projection of related individuals (if desired), generation of PCA figures that assist in setting the best number of principal components (PCs) for association analysis, availability of mixed models, use of recommended software for GWAS based on population size, and a Markov chain Monte Carlo (MCMC) method to estimate best set of PRS parameters. Finally, we tested GARSA pipeline in a family-based Brazilian admixed population and demonstrated that the corrections implemented indeed mitigate bias in downstream analysis. The pipeline can be implemented on personal or server-side environments.

**Availability:** The development version (open-source) is available in https://github.com/LGCM-OpenSource/GARSA

**Contact:** Fernando P. N. Rossi -; José S. L. Patané -

**Supplementary information:** Supplementary tutorial.

## Introduction

The completion of the human genome enabled the creation of human genetic variant databases, which in conjunction with the evolution of genotyping technologies allowed the first Genome-wide association studies (GWAS) (The Wellcome Trust Case Control Consortium, 2007). GWAS is a first step to identify candidate variants associated with the genetic mechanisms underlying complex phenotypes in humans. Briefly, it is based on a regression model, where the association between thousands to million genetic variants, either single nucleotide polymorphisms (SNPs) or insertion/deletion sites (indels), is tested against a given phenotype. The model also includes covariates known to influence the studied phenotype, such as age, sex, and principal components. For human complex phenotypes and common diseases, GWAS has been instrumental in expanding the number of candidate genomic loci relevant to the biomedical area (Visscher et al., 2017).

A Polygenic risk score (PRS) estimates an individualized index of the aggregated influence of all genetic variants tested in a GWAS against a phenotype, and can be used as a predictor of the development of an outcome (e.g., death, hypertension). Specifically, it is a renormalization of the effect sizes of each of the GWAS variant regressors while also taking into account the number of alleles per individual.

Together, these two analytical tools are evolving and becoming instrumental to understand the genetic architecture of complex phenotypes/diseases. Here, we present a Python-based semi-automated pipeline, which unifies the steps, and widely used softwares, to implement these analyses, from preprocessing the dataset for GWAS, up to PRS. We also included options targeted at processing data from admixed populations, expanding most of the studies that are focused on European-based (and generally non-admixed) populations.

## Description of the GARSA pipeline

### Functionalities

The overall workflow of Genomic Association Risk Score for Admixed populations (GARSA) is divided into nine modules (Supplementary tutorial) as depicted in Fig. 1 and described below, with special focus on the Kinship and PCA modules. In most modules, the datasets are provided as VCF or PLINK (https://github.com/chrchang/plink-ng)binary files (.bed, .bim, .fam). At each step, the output receives a new extension, allowing the user to easily navigate through the outputs. This is important, since the pipeline does not demand a fixed order for the use of each module, and therefore these can be used in any order desired.

**Fig. 1:**
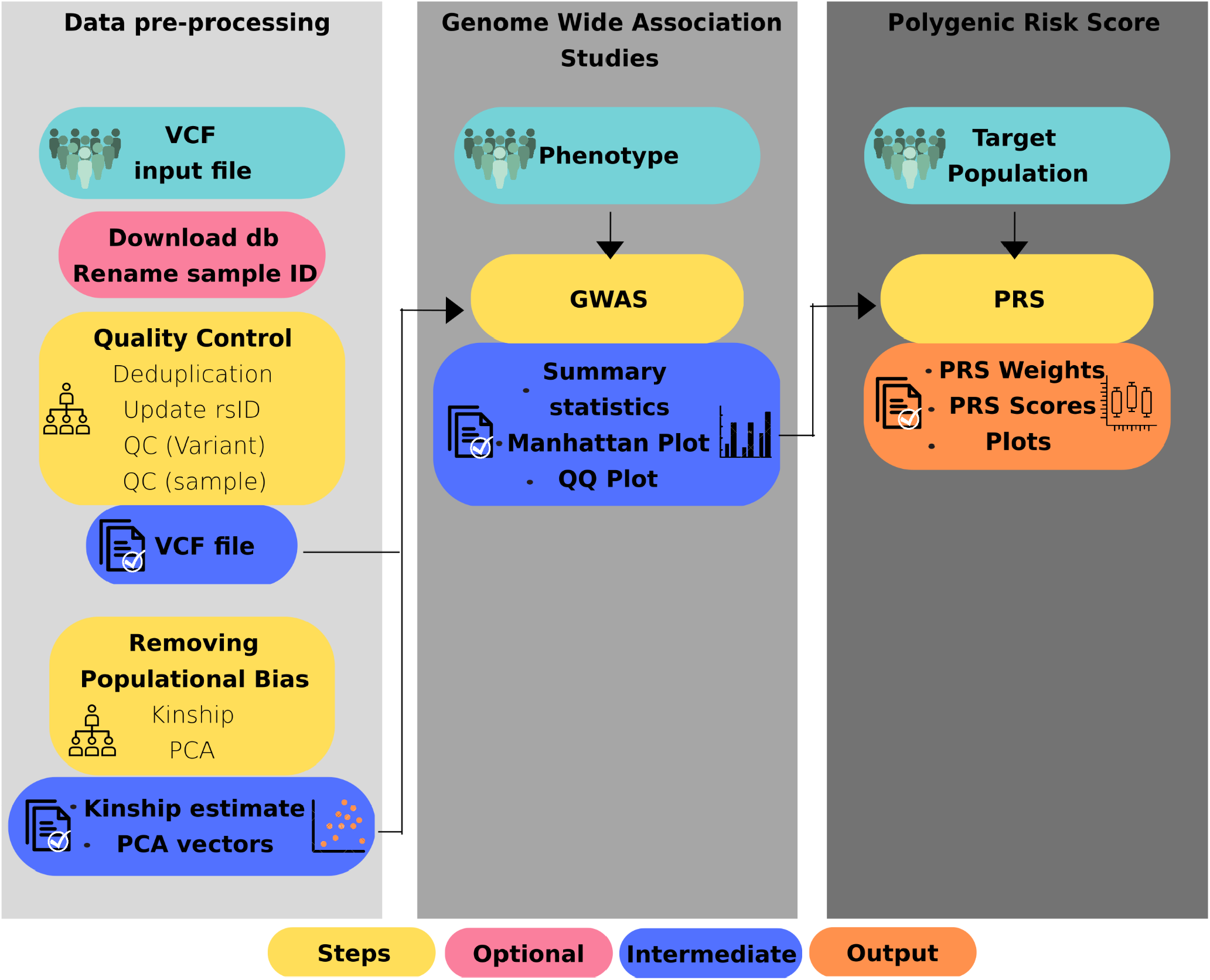
Functionalities and associated modules within the GARSA framework.

Aiming at facilitating data manipulation on each step of the processing pipeline and proper handling in every module that uses PLINK (both 1.9 and 2.0), we recommend the formatting of every sample ID to be “Family ID” (FID) followed by “Individual ID” (IID), separated by an underscore. If your dataset lacks a Family ID, the Individual IDs (e.g. IID_IID) can be duplicated. To aid in formatting data, we provide a module called “rename_sample_id” that takes as an input a tabular (or comma) delimited file containing the original sample name in the first column, and in the second column the new sample name formatted as FID_IID.

### Deduplication/multiallelic removal module

This module uses PLINK binary files, so when a VCF is provided, it is automatically converted into that format. After conversion, all variants are loaded and searched for multiallelic variants using PLINK 1.9, and duplicated variants using bash scripts. In both cases, those variants are removed from subsequent analyses.

### Annotation of rsIDs

With the output file from the module above, an annotation step can be performed using the “update_rsID” module. In this step, the pipeline includes the rsID for each variant considering its chromosome, position, reference and alternative allele, using bcftools (v. 1.10.2)*annotate* function and dbSNP (version b151, downloaded using the download_db script) (Day, 2010) reference file. To allow the annotation of the largest possible number of variants, it also includes swapping (which allele is considered reference, and which one is the alternate allele) and flipping (of the DNA strand) corrections of variants in the target population. It works by considering a human genome reference VCF file (hg37 or hg38) parsed into a table format, which gets compared to the user’s VCF using python scripts, allowing allele flippings and swappings to be identified and corrected using Plink.

### Variant and sample quality control

The module named “quality_control” applies variant quality control from PLINK2, mostly using guidelines from Anderson et al. (2010), to remove poorly imputed data (R2 and INFO score), variants with high missing genotype data and with minor allele frequency (MAF) below a given threshold, and filters variants using a Hardy-Weinberg equilibrium (HWE) threshold. The HWE flag can be either turned on or off depending on the user’s preference; default is off since this analysis can interfere with the population structure (Pearman et. al, 2022). The module receives as input a VCF file, and allows customization of parameters and thresholds.

For sample quality control, we developed a module named “quality_ind”, which uses PLINK1.9 to filter samples with high missing data (“mind”) and high heterozygosity rate (HET) and remove them from the dataset. To achieve better performance, we recommend the concatenation of all chromosomes into a single VCF file.

### Kinship and PCA analysis

The modules for Kinship and PCA analyses are responsible for the minimization of kinship- and population-related biases, using approaches best suited for highly admixed populations.

For the Kinship module, we used the approach proposed for Conomos et. al (2016) to circumvent the admixture levels present in many current populations (e.g. Brazilian) and hence correct the Kinship values based on population structure. This step is based on R libraries SNPRelate, GENESIS, SeqArray, and SeqVarTools. The correction uses PCA vectors as covariates to correct previously estimated kinship values, reducing population structure bias and the bias introduced by negative kinship values in the dataset.

For the PCA module, we used the FlashPCA program, whose output is processed by our scripts to detect outlier variant loads that might confound the population structure with loadings that may be specific to genotype structure. In this way, we minimize the noise caused by outlier variants that may bias population structure. The analysis is based on the R library BigSNPR.

### Genome Wide Association (GWAS) and Polygenic Risk Score (PRS)

The GWAS and PRS modules are the last two modules in the pipeline. In the GWAS module, the user can choose between two regression tools based on the dataset sample size: GCTA (Yang et al., 2011) or BOLT-LMM (Loh et al., 2018). The GCTA tool is best suited for smaller datasets (hundreds of samples) and BOLT-LMM for large datasets (best if larger than 5,000 samples). A choice can be made for correcting kinship analysis for GCTA (BOLT-LMM uses its own kinship matrix estimate). Also, in both of those modules the user must provide a formatted table (described in the documentation) containing phenotype information for every sample being processed in the analysis.

For the PRS module, the GWAS output can be used alongside the same phenotype information used before. In this step, the PRS for each sample will be calculated using LDPred2 (Privé et al., 2020) and plotted using a python script. The results from both modules can be presented as figures and tables.

### Application in admixed dataset

To demonstrate the importance of Kinship and PCA bias correction in GWAS and, consequently, PRS analysis, we used GARSA to apply QC and GWAS regression in an admixed Brazilian dataset of 2,048 individuals, with strong family structure. We compared the results of the GARSA pipeline (in the kinship and PCA module) with the King robust implementation in Plink2 (--make-king) and FlashPCA without loading correction. For both strategies, the resulting data (PCA vectors and kinship matrix) were used as covariates for GWAS regression analysis for height phenotype, using GCTA Linear Mixed Model (LMM).

We observed relevant differences on the kinship estimate from both strategies tested, with a higher number of negative kinship values estimated by Plink (Fig. 2A). Kinship negative values are difficult to interpret and can introduce bias on the downstream analysis (Jiang et al., 2022). Therefore, we demonstrated that GARSA can mitigate this bias, specially in admixed population. Also, we observed a significant discrepancy between kinship values between two individuals estimated by both tools (Fig. 2B), for example, where GARSA estimates individual relationships as second degree, while King robust estimates as first or third degree (left panel of Figure 2B). Interestingly, when we checked the relationship between individuals in the database, considering the pedigrees, we observed that GARSA correctly estimated first degree individuals in 92% of the cases, while Plink correctly estimated only 66%.

**Fig. 2:**
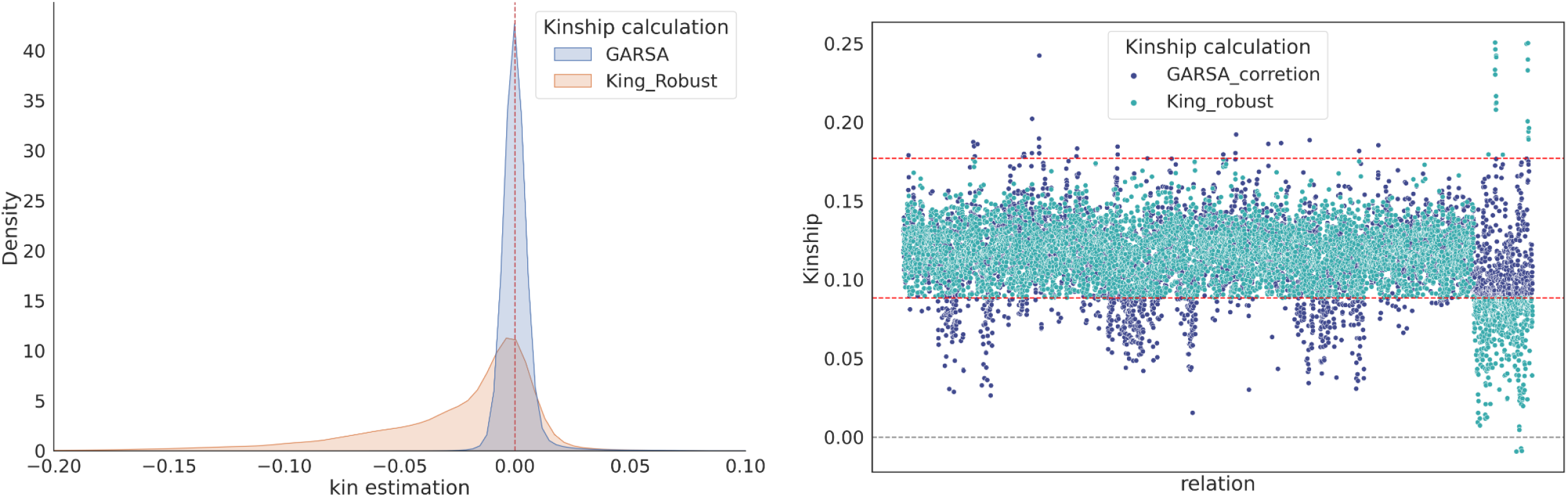
(**A**) Distribution of kinship estimates for GARSA (blue) and King Robust (orange). The line in red marks the point of “zero” relatedness. (**B**) Differences in the Kinship estimate for both strategies, in green the King Robust and in blue GARSA. The red dashed lines mark the limits for second degree estimation, and gray lines (at 0.0) indicate no relationship.

We further observed differences in the amount of significant variants detected on the GWAS results from both strategies, identifying 144 significant variants for height using kinship and PCA correction by GARSA and only 87 significant variants using kinship and PCA estimates by Plink. The difference in the number of significant variants between the analyzes is also observed in the number of genomic risk loci for height using FUMA web application. Considering GWAS using GARSA pipeline, we identified 12 risk loci, while only 5 risk loci were identified using the Plink strategy, with a single risk locus present in both analyses. These results demonstrate the importance of bias correction in GWAS analysis to improve detection of relevant loci, specially in family-based admixed populations.

## Discussion

GARSA is a Python semi-automated and ready-to-use pipeline, which applies recommended practices for GWAS and PRS analysis, including options that take into account population structure for dealing with admixed populations that otherwise bias the overall results.

Regarding GWAS, multiple preprocessing steps are used to correct for kinship while also removing populational confounders; identification and removal of outlier variants with extreme loadings are performed; allows Principal Components re-projection for related individuals, avoiding loss of samples; plots first PC against higher-order ones, to facilitate determination of the most reliable number of PCs to be use as covariates in the GWAS analysis; allows the choice of the software to be used, depending on the dataset size; and provides a linear mixed model that allows regression analysis considering related individuals. Regarding PRS, the pipeline implements a Markov chain Monte Carlo (MCMC) method to estimate the best set of PRS parameters.

As a proof of concept, we applied the pipeline in a family-based admixed population with 2,048 individuals and 7.8 million variants. The results showed a significant improvement in the number of both genomic risk loci and significantly associated variants in GWAS, and also in the removal of relatedness bias, when compared to the “standard” approach using PLINK.

## Supporting information

Supplementary Table 1: Genomic Risk Loci identified in GWA studies for height in admixed Brazilian population, using Plink and GARSA

## Acknowledgements

We thank Renata Callado and Marco Antônio Gutierrez for the bioinformatic infrastructure and management, and Manuela Bonetto for the additional code review. We also thank the participants and investigators of UK Biobank, Baependi Heart Study, and EpiGen-Pelotas for allowing pipeline testing in their datasets.

